# A Systematic Review of the Impacts of El Niño-Driven Drought, Fire, and Smoke on Non-Human Primates in Southeast Asia

**DOI:** 10.1101/2024.11.08.622697

**Authors:** Elizabeth J. Barrow, Wendy M. Erb

## Abstract

During the El Niño phase of the El Niño-Southern Oscillation (ENSO), much of Southeast Asia experiences intense droughts and surging temperatures, exacerbating wildfires and creating a blanket of hazardous haze across much of the region. These patterns are predicted to worsen with climate change, further exacerbating the vulnerability of the regions’ primate species, 94% of which are threatened with extinction. Here, we report findings from a systematic search of the literature and synthesise the current state of knowledge regarding the impacts of El Niño-driven heat, drought, fire, and smoke on non-human primates in Southeast Asia. Our review shows that habitat loss and degradation driven by El Niño-induced drought and fire causes changes to diet, activity patterns, social interactions, and physical condition, and leads to population crowding, reduced group sizes, increased infant mortality, displacement of individuals, and local extirpation. Further, prolonged exposure to smoke alters behaviour and worsens the health of primates. Notably, two studies presented evidence for the recovery of primate populations in habitats damaged by fire if forest is allowed to regenerate. We highlight significant gaps in our understanding of the impacts of El Niño on non-human primates, particularly the need for research that encompasses the interconnected factors of rainfall, temperature, fire, and smoke, as well as long-term effects on reproduction and mortality. Due to the unpredictable nature of El Niño, we acknowledge the difficulties of planning research in this area and emphasise the potential of long-term research sites to add to our understanding of this key conservation issue.

## Introduction

The El Niño-Southern Oscillation (ENSO) is a natural climate phenomenon that changes both atmospheric and oceanic oscillation (Niedzielski, 2014). ENSO has three phases defined by changes in sea surface temperature and trade winds over the central and eastern Pacific Ocean (Niedzielski, 2014): El Niño (warm phase), La Niña (cold phase), and neutral (neither El Niño nor La Niña). The cold and warm phases have significant influence on environmental processes, most notably causing changes to weather patterns - including increased rainfall and flooding in some regions, and decreased rainfall and drought in others (Zhou et al., 2014).

The El Niño phase of ENSO is characterised by above-average sea surface temperatures in the central and eastern tropical Pacific Ocean, during which a weakening of wind speed or change in direction can cause higher-than-average rainfall over South America and drought across Southeast Asia (Climate Prediction Center Internet Team, 2012). The onset, progression, and persistence of El Niño conditions vary, but they typically develop during the Northern Hemisphere spring or summer and often peak in winter (Liu et al., 2025). El Niño fluctuates in frequency and intensity, occurring approximately every 2-7 years (Wang et al., 2019), with events defined as moderate or strong (Santoso et al., 2017; Wang et al., 2019). A moderate El Niño event is defined as when sea surface temperatures are 0.5-2.5 °C above average. When temperatures exceed 2.5 °C the event is classified as a strong El Niño (Wang et al., 2019). Strong El Niños have caused catastrophic natural disasters worldwide, including flooding (Vos, 1999), drought (Lestari et al., 2018), and wildfires (Khandekar et al., 2000).

While the overall frequency of El Niño has remained relatively stable over time, there has been a sharp increase in the occurrence of *strong* events, pointing to a rising trend in the intensity of this phenomenon (Wang et al., 2019). During the 60-year period between 1901 and 1960, just one strong El Niño event occurred, but the subsequent 60-year period (1961-2020) experienced four (Wang et al., 2019; Climate Prediction Center Internet Team, 2024). Oceanic warming is likely responsible for the increasing severity of El Niño events, a trend that is predicted to worsen with the changing climate (Thirumalai et al., 2017; Wang et al., 2019).

In the western equatorial Pacific, particularly in Southeast Asia, El Niño causes reduced rainfall and increased temperatures (Thirumalai et al., 2017; McAlpine et al., 2018; Rifai et al., 2019; Fig. 2). However, the effects of El Niño are not uniform across the region. The southern part of Southeast Asia (below 5°N) experiences stronger effects of El Niño than the northern part, primarily due to ocean surface currents in that area and proximity to the equator (Hariadi et al., 2021). During 2015’s strong El Niño event, Southeast Asia experienced record-breaking high temperatures and low rainfall, particularly in Peninsular Malaysia, southeastern Borneo, and southern Papua New Guinea (Rifai et al., 2019). These extremes are thought to be driven by climate change, highlighting the region’s increasing vulnerability to future events (Thirumalai et al., 2017; Rifai et al., 2019).

**FIG. 1:**
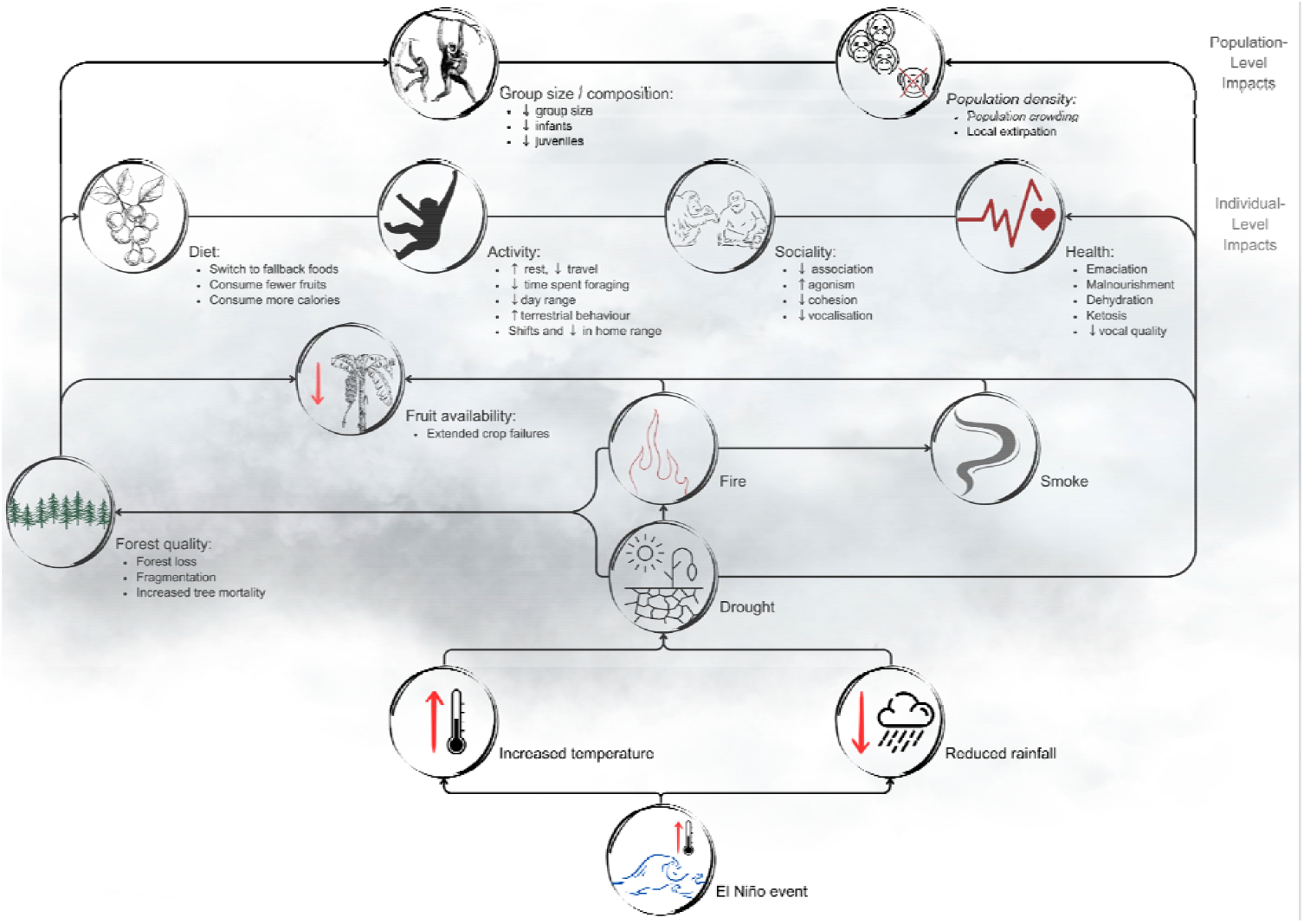
Conceptual diagram illustrating the negative impacts of El Niño on Southeast Asian primates based on our systematic literature review.

**FIG. 2:**
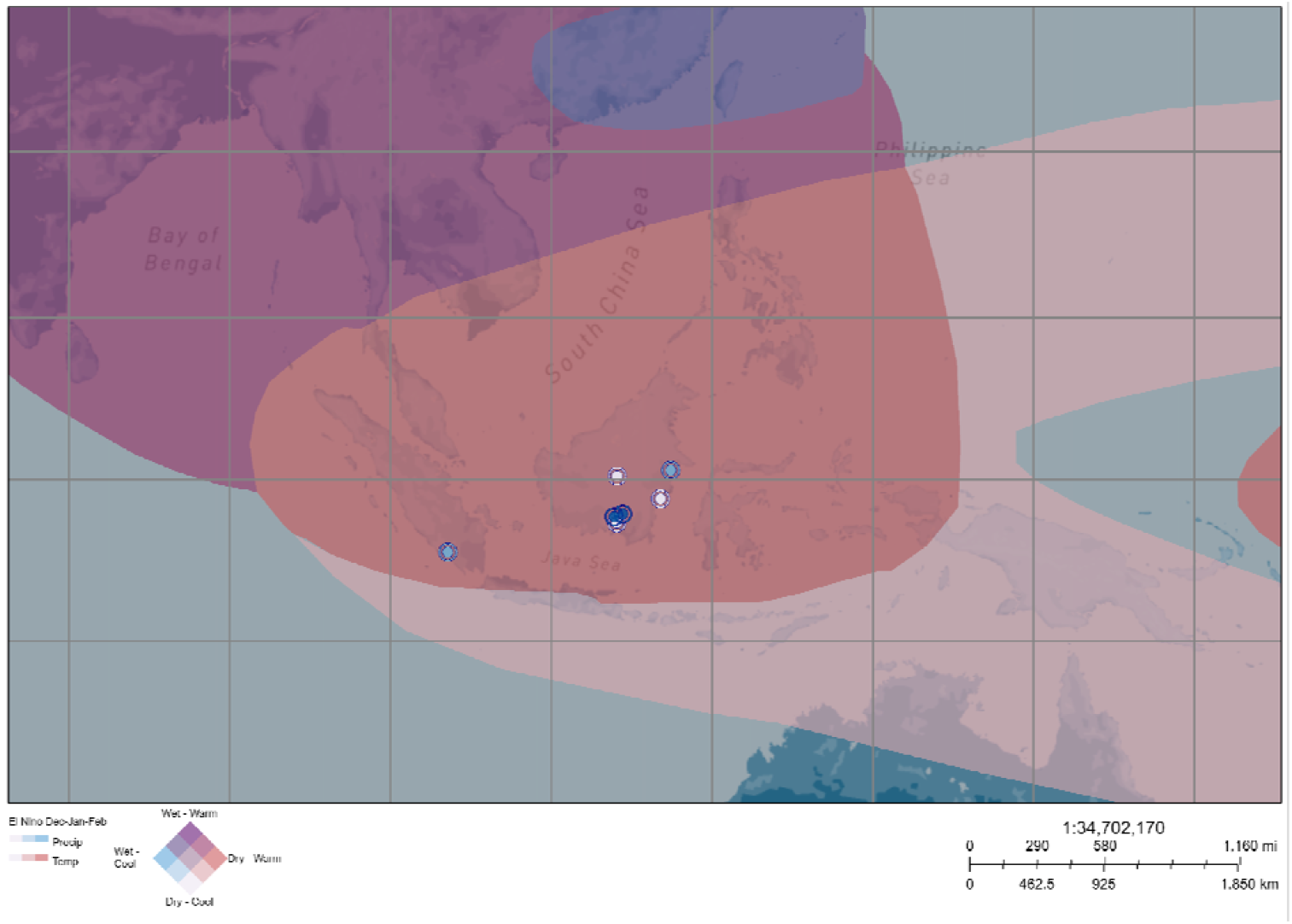
Map of El Niño’s weather effects from December to February across Southeast Asia. Location of study sites included in our review are indicated by blue circles (darkness indicates number of publications at each site, ranging from 1 = lightest to 6 = darkest). Map created using “ENSO Global Weather Patterns” by joe.riedl_portlandcc, hosted on ArcGIS Online (Esri, 2021).

The compounding effect of reduced rainfall and elevated temperatures significantly contributes to more frequent, intense, and widespread wildfires (Wooster et al., 2012; Page & Hooijer, 2016), which causes widespread hazardous haze (Marlier et al., 2013; Ramakreshnan et al., 2018; Shokoohinia et al., 2020). This is particularly significant in areas with extensive peatlands, such as Southeast Asia, which holds over half of the world’s tropical peatlands (Page et al., 2011). Although peatlands are naturally fire-resistant, rapid degradation caused by deforestation and drainage for agricultural conversion, particularly in southern Southeast Asia (e.g., Indonesia, Malaysia), has dramatically increased their susceptibility to fires (Page et al., 2011; Page & Hooijer, 2016; Harrison et al., 2024). Repeated fires in these areas further degrade habitats, making them more vulnerable to long-term damage, including reduced resilience and loss of biodiversity (Cochrane, 2003; Page & Hooijer, 2016). In Indonesia and some parts of Malaysia, fires are frequently used to clear land. While intended as controlled burns, these fires often spread uncontrollably, especially during dry seasons intensified by El Niño events, leading to severe wildfires with extensive environmental and health consequences (Khan et al., 2020).

El Niño-induced extreme weather patterns have far-reaching impacts on human health and wellbeing. Drought can lead to catastrophic damage to farmlands, with devastating consequences for agriculture. Between 2015 and 2019, over 17 tonnes of crops were lost as a result of drought, impacting the lives of over 100 million people in Southeast Asia (Venkatappa et al., 2021). During severe fire seasons, typically associated with strong El Niño events, widespread haze significantly reduces air quality, extending as far as southern Thailand (Shokoohinia et al., 2020). This haze has been linked to human deaths in Indonesia, Malaysia, and Singapore (Roy & Goh, 2019), and is estimated to cause the premature death of 33,100 adults and 2,900 infants annually on Borneo and Sumatra (Hein et al., 2022). Furthermore, it has led to school and airport closures, reduced job opportunities, and disrupted transportation of goods, impacting local quality of life and national economy (Limin et al., 2006). Biological communities in Southeast Asia are also impacted by El Niño, but the impacts are not always uniform across ecosystems. Whereas drought conditions can cause increased fruit availability in dipterocarp forests in East Kalimantan by triggering masting (i.e., simultaneous mass fruiting of many species) (Fredriksson et al., 2006); in Central Kalimantan’s peat swamp forests, the combination of drought, heat, fire, and smoke can result in an extended period of lower-than-average fruit production (Ashbury et al., 2022). More broadly, a decline in photosynthesis was observed in Southeast Asia during the strong 2014-16 El Niño event, suggesting that drought conditions generally reduce plant productivity (Qian et al., 2019). El Niño-driven fires and haze on Borneo have also been linked to increased litter-fall (Harrison et al., 2007), reduced photosynthesis (Davies & Unam, 1999), increased tree mortality (Berenstain, 1986; O’Brien et al., 2003; Russon et al., 2015), and reduced biodiversity (Harrison et al., 2024). Haze has far-reaching impacts on biodiversity, exemplified by a Singapore-based study that demonstrated a drastic decline in acoustic activity due to haze from a strong El Niño event in 2015, originating from forest fires on Borneo and Sumatra (Lee et al., 2017).

Southeast Asia faces ongoing environmental challenges that threaten its biodiversity, which are further exacerbated by El Niño. The region is experiencing the highest rates of deforestation of any tropical area, leading to the loss of biodiversity at an unprecedented rate (Sodhi et al., 2010). Currently, more than 3,000 of the region’s plant and 2,500 animal species are threatened with extinction (IUCN, 2021). Due to the high concentrations of endemic species, loss of populations in this world-renowned biodiversity hotspot (Myers et al., 2000) will likely result in global extinctions (Sodhi et al., 2004).

Southeast Asia’s primates face a particularly dire situation, with a staggering 94% of species threatened with extinction, far outpacing the global average for primates (Estrada et al., 2017; IUCN, 2021). Their primary threat is habitat loss due to agricultural expansion, logging, livestock farming, poaching, and other forms of removal from the wild (Estrada et al., 2017), with climate change increasingly recognised as a factor in their decline (Graham et al., 2016; Estrada et al., 2017; Zhang et al., 2019; Bernard & Marshall, 2020). More extreme weather patterns are likely to render more habitat unsuitable and disrupt food availability (Graham et al., 2016; Estrada et al., 2017; Bernard & Marshall, 2020). As the impacts of El Niño are predicted to worsen with climate change (Thirumalai et al., 2017), it is probable that the climate-related impacts faced by primates may be even more pronounced in areas worst affected by El Niño.

In this review, we focus on how El Niño impacts primates inhabiting Southeast Asia. Despite the growing recognition of climate change as a significant factor threatening primates, research in this area has only become a topic of interest within the last 25 years and is minimal compared to other taxonomic groups (Bernard & Marshall, 2020). The regions where primates live are expected to experience 10% more warming than the global average, highlighting their heightened vulnerability to climate-related changes (Graham et al., 2016). Southeast Asia is expected to experience particularly severe climate impacts, with El Niño events intensifying and endangered species among the most vulnerable to these changes (Graham et al., 2016; Wang et al., 2019). Understanding how these changes affect primates is urgent, given their high conservation risk and critical role in contributing to ecosystem health and function (Estrada et al., 2017). By synthesising current research into the impacts of El Niño-driven heat, drought, fire, and smoke on Southeast Asian primates, our review aims to (1) assess the current state of knowledge, (2) highlight gaps in our knowledge, and (3) identify opportunities for future research. Combined, this information will provide crucial guidance for future research and conservation initiatives in Southeast Asia.

## Methods

On April 30, 2025, we conducted a systematic review of English-language literature using Web of Science. We constructed three sets of search terms to encompass study region, taxa, and topic. For study region, we specified all 11 Southeast Asian countries, including alternate names (e.g., both Myanmar and Burma were included). To build a complete list of taxonomic terms, we searched the IUCN Red List (IUCN, 2024) for all primate species known to inhabit these 11 countries. Using this list, we included as search terms both common names and genera. To search by topic, we included ENSO (and alternate names, e.g., El Niño) along with El Niño-related factors (i.e., temperature, rainfall, drought, fire, smoke, haze, and smog). Search terms *within* each of the three sets were separated using the boolean operator “OR” and were then combined *across* these three sets using the boolean operator “AND” to create a single search. We entered all search terms under the category “Topic”, which searches for terms in the title, abstract, and keyword fields, and performed the search using “All Databases” (Supplementary Fig. 1 and Supplementary Material 1).

After removing duplicates (*n* = 15 of 376), we reviewed the results by article type, ensuring that only peer-reviewed journal articles were included for further review (*n* = 257). Next, we screened the articles by title, and only those mentioning any of the species of interest, El Niño or El Niño-related factors, or one or more of the included countries were kept for the next stage (*n* = 88). We then reviewed abstracts, only including studies of primates in the region of interest that mentioned El Niño, drought, fire, smoke, temperature, or rainfall (*n* = 32). We then conducted a full-text review of the remaining articles. Lastly, we included studies that explicitly mentioned El Niño as a contributing factor to reduced rainfall, increased temperature, fire, or smoke. Additionally, we included studies focusing on drought, fire, or smoke that overlapped substantially with one or more El Niño events, even if El Niño was not explicitly mentioned, due to the stronger and more direct links between these phenomena and extreme climate events (Supplementary Table 1). We summarised the final 15 studies by location, taxon, El Niño event(s), and El Niño-related factor(s) (Table 1).

**TABLE 1:**
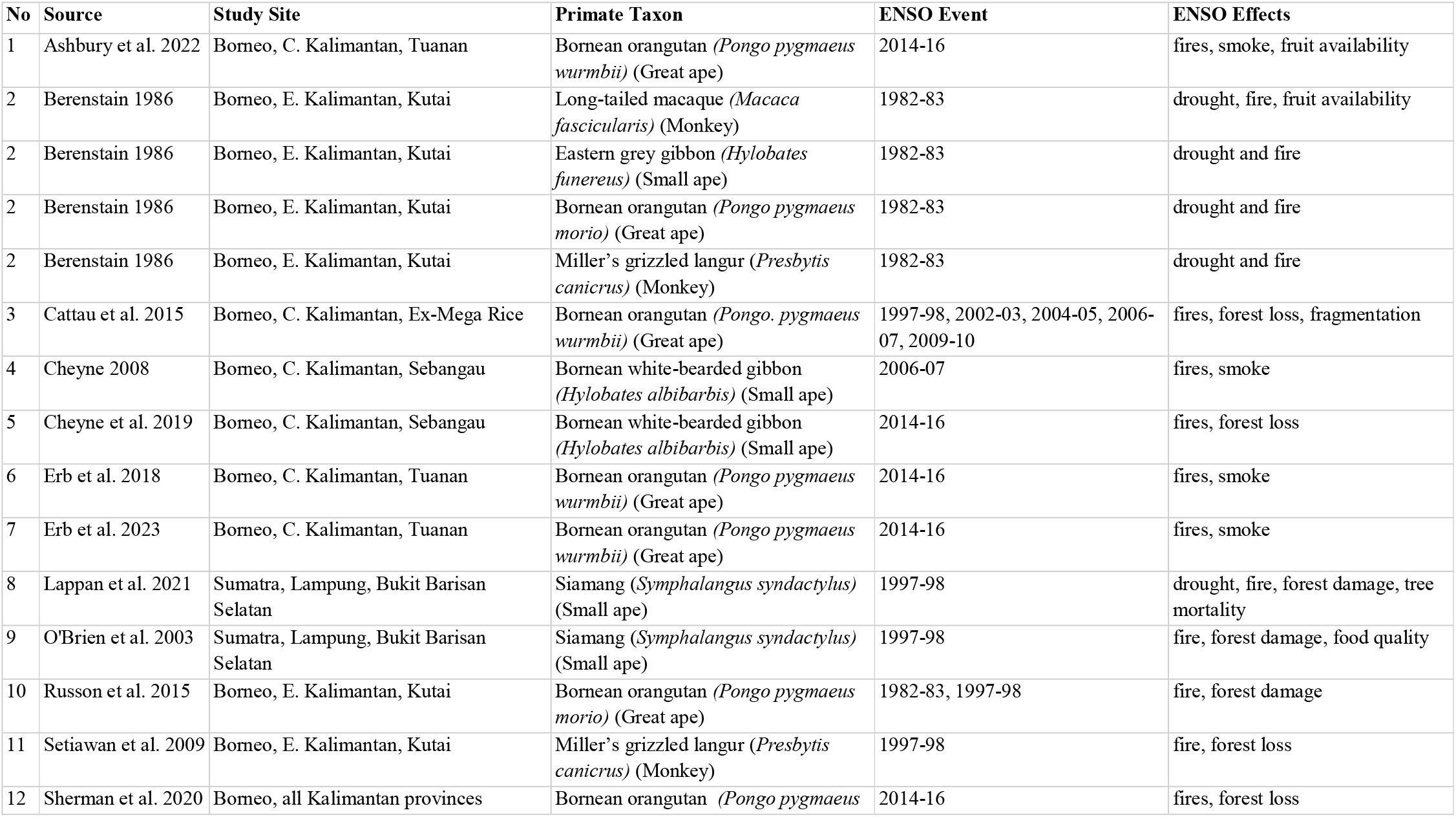
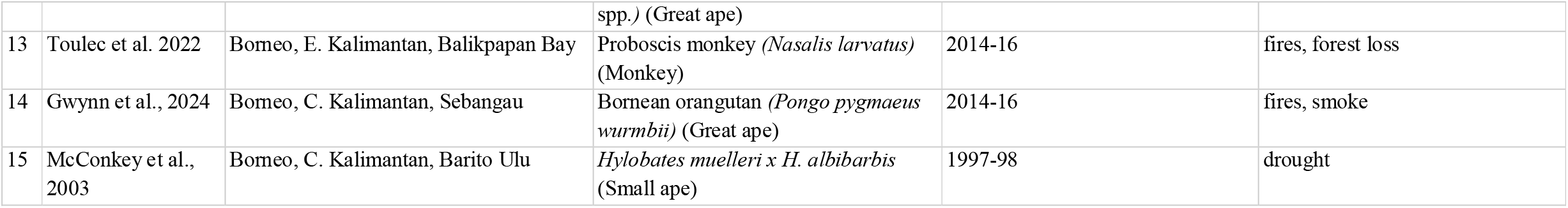
List of studies included in our review. For each entry, we provide the island, province, and site where the study took place (all were conducted in Indonesia), taxon, date(s) of El Niño, and the El Niño-related effects. Where studies examined numerous species, they are listed more than once.

### Geographic And Taxonomic Representation

The 15 articles demonstrate limited geographic and taxonomic representation of research on El Niño-driven environmental impacts on primates within Southeast Asia. All of the studies were conducted in Indonesia. Among these, 86.7% (13 of 15) were conducted in Borneo (Kalimantan) and the remainder in Sumatra. Of the studies in Borneo, 61.5% (8 of 13) were from three sites in Central Kalimantan, 30.8% (4 of 13) from two sites in East Kalimantan, and 7.7% (1 of 13) covered all Kalimantan provinces. In Sumatra, both studies were from the same research site in southern Sumatra (Table 1).

Taxonomically, studies focused on diurnal apes and monkeys, with no representation of the region’s nocturnal tarsiers or lorises (Table 1). The largest proportion (53.3%, 8 of 15) examined the Bornean orangutan (a great ape), covering all three subspecies: *Pongo pygmaeus wurmbii, P. p. morio*, and *P. p. pygmaeus*. Small apes were the subject of 40% of studies (6 of 15), including Sumatra’s siamang (*Symphalangus syndactylus*) and Borneo’s white-bearded (*Hylobates albibarbis*), northern gray (*Hylobates funereus*), and *Hylobates muelleri x H. albibarbis* (previously known as *Hylobates muelleri x H. agilis*) gibbons. Borneo’s monkeys were the subject of 20% of studies (3 of 15), covering the long-tailed macaque (*Macaca fascicularis*), Miller’s grizzled langur (*Presbytis canicrus*, previously known as *P. hosei*), and the proboscis monkey (*Nasalis larvatus*). The majority of studies in our review focused on a single species, but one covered multiple sympatric primates.

### El Niño Events and Effects

The articles in our review investigated El Niño events from 1982 onwards, with a predominant focus on strong events, including 1982-83 (13.3% of studies, 2 of 15), 1997-98 (40%, 6 of 15), and 2014-16 (46.7%, 7 of 15) (Table 1). Only one study specifically examined the impacts of a moderate El Niño event (2006-07). Most studies (73.3%, 11 of 15) concentrated on a single El Niño event, while the remainder (26.7%, 4 of 15) utilised longer datasets overlapping multiple events, or examined impacts in areas previously affected by more than one El Niño event. The majority of articles (53.3%, 8 of 15) examined the direct impact of habitat loss due to fire. Three others (20%) investigated the impacts of reduced habitat quality (including changes in food availability) resulting from drought, fire and/or smoke, while the remaining four articles (26.7%) focused specifically on smoke (Table 1).

The studies took diverse approaches to investigate the impacts of El Niño-driven drought, fire, and smoke on primates. Nine studies conducted research during an El Niño event, but also collected data over a longer period of time, or utilised long-term datasets before and/or after the studied environmental factors (e.g., fire, smoke, or drought) to investigate changes and recovery over time (Berenstain, 1986; Mcconkey et al., 2003; Cheyne, 2008; Russon et al., 2015; Erb et al., 2018; Cheyne et al., 2019; Ashbury et al., 2022; Erb et al., 2023; Gwynn et al., 2024). These datasets varied significantly in length, spanning from 6 months to over 40 years, presenting evidence for both short- and long-term impacts. Two studies conducted research immediately after El Niño-induced fires and continued long-term data collection for 3 years in one case (O’Brien et al., 2003) and 18 years in the other (Lappan et al., 2021). Three articles involved population surveys conducted in areas directly affected by and following an El Niño event, but the data collection period did not directly overlap the event. Two of these were singular surveys conducted after one or more El Niño event (Setiawan et al., 2009; Cattau et al., 2015), while the third compared three separate surveys covering a 10-year period, before and after the 2014-16 strong El Niño event (Toulec et al., 2022). One final study documented a sharp increase in translocations of Bornean orangutans during the 2015 El Niño-driven fires, highlighting the direct impact of habitat loss (Sherman et al., 2020).

### Impacts on Primates

Among the 15 articles in our review, three major themes emerged concerning the impacts of El Niño on primates, including 1) population size and structure, 2) behaviour (diet, activity, ranging, and social interactions), and 3) health and physical condition.

#### Population size and structure

Six studies examined the effects of fires on populations of primates on Borneo and Sumatra (Table 1). In the four months following the 1983 fires in Kutai National Park (Borneo), Berenstain (1986) observed groups of long-tailed macaques, orangutans, gibbons, and Miller’s grizzled langurs. During this period, he documented no loss of groups or mortality for any of these species. Similarly, two years after the 2015 fires, a survey of proboscis monkeys in Balikpapan Bay (Borneo) showed no signs of decline (Toulec et al., 2022).

In contrast, the four studies that examined the impacts of the 1997 fires documented significant changes in primate populations. Two studies, conducted a decade after the 1997 fires, found dramatic changes in Bornean primate populations. Ten years after the fires in Kutai National Park, surveys of Miller’s grizzled langurs found an alarming disappearance of the species from this site (Setiawan et al., 2009). Surveys in the Ex-Mega Rice Project area found that orangutan population density had dramatically increased due to crowding in small forest fragments that were created by fires 12 years prior to the study (Cattau et al., 2015).

In the years after the 1997 fires, demographic changes in groups of siamangs at Bukit Barisan Selatan National Park (Sumatra) were documented over two different time periods. Three years after the fires, O’Brien et al. (2003) documented that groups inhabiting burned areas had a higher proportion of adults and subadults compared to groups in non-burned areas (74% vs. 63%) and also showed lower survival of infants and juveniles. Eleven years after the fires, (Lappan et al., 2021) found that groups whose home ranges overlapped burned areas were still smaller than those that ranged in unburned areas.

#### Behaviour

Five studies examined the effects of drought, fire, and smoke on the diets and activity patterns of long-tailed macaques, gibbons, and orangutans in Borneo. Before, during, and after the 1983 fires Berenstain (1986) made full-day behavioural observations of one group of long-tailed macaques in Kutai National Park. He documented a sharp increase in the proportion of time the monkeys spent feeding on insects/caterpillars, foliage, stems, petioles, and unripe seeds of dipterocarps in 4 months after the fires. During this time, the macaques also increased resting, and decreased foraging time.

McConkey et al. (2003) studied food choice of two groups of hybrid gibbon groups before and during a severe drought in 1997 in Barito Ulu, Central Kalimantan. They showed that gibbon food choice was influenced by weather-driven shifts in food availability. When fruit was scarce during periods of low rainfall, gibbons consumed more flowers, figs, and young leaves. An unusual peak in flower consumption was recorded during the drought, which was linked to an El Niño-induced mast that caused an atypical abundance of flowers.

Russon et al. (2015) studied orangutan feeding, diet, and activity patterns at Kutai National Park following the 1997 drought and fires. They showed that, in the first year after the fires, orangutans ate twice the number of top foods (i.e., plant taxa representing >20% of feeding time) and consumed overall a more diverse diet (50-100% more species) comprising relatively few fruit foods. By 12-15 years post-fire, however, the diet had recovered to original patterns. In the 3-4 years post-fire, orangutans spent less time feeding and more time resting; but had shifted toward pre-fire activity patterns after 12-15 years, corresponding with changes in the availability and distribution of food.

Two studies examined the effects of smoke from the 2015 fires on adult female and flanged male orangutans at Tuanan. Ashbury et al. (2022) compared activity and diet of adult females between 2010-2018 during low-fruit periods before and (up to 32 months) after the fires. They showed that female orangutans adopted energy-conserving shifts in their diet and activity patterns post-fire. In particular, in the low-fruit periods following the fires, females rested more and travelled less, while switching to less-preferred foods (e.g., bark, pith, mature leaves, etc.). In another study, Erb et al. (2018) compared activity and diet of adult flanged males for 5 months before, 2 months during, and 3 months after the 80-day period of poor air quality caused by the 2015 fires. In this study, adult flanged males rested more during the smoke and post-smoke periods when compared to the pre-smoke baseline. Similarly, during the post-smoke period, they travelled less while simultaneously consuming more calories compared to pre-smoke levels.

The impacts of fire and smoke on primate ranging and group cohesion were documented in four studies of macaques, gibbons, and orangutans in Borneo, and siamangs in Sumatra. In the four months following the 1983 fires in Kutai National Park (Borneo), long-tailed macaques greatly reduced their day range, exhibited more terrestrial behaviour, and reduced group cohesion (Berenstain, 1986). In Bukit Barisan Selatan National Park (Sumatra), O’Brien et al. (2003) and Lappan et al. (2021) studied the home ranges of seven siamang groups following the 1997 fires. Groups whose home ranges included fire-damaged habitat had not substantially expanded their ranges into the recovering area during the 3–11-year period following the fires. By 18 years post-fire, however, two of the three groups’ home ranges included substantial areas of burned habitat. Similarly, in 2002 (5 years after the fires), no siamang groups were observed in the burned area interior, but between 2012 and 2017, 2 groups and 2 or more solitary individuals were observed ranging there.

Two studies documented changes in ranging behaviour of apes in Borneo after the 2015 fires. In Sebangau Forest, Cheyne et al. (2019) documented a shift in home range location for the group of white-bearded gibbons whose range was damaged by fire (up to 3 years post-fire). For the three adjacent groups whose ranges did not overlap burned habitat, no changes were observed in their home range locations. At Tuanan, flanged male orangutans’ daily path length decreased in the 3 months following the smoke compared to the 5-month pre-smoke period (Erb et al., 2018).

Lastly, changes in social behaviour in response to wildfire smoke were documented in three studies of apes on Borneo. White-bearded gibbons in Sebangau reduced the number of singing days and song bout length on smoky days during the 2006 fires compared to non-smoky days (Cheyne, 2008). In the 3 months following the 2015 fires, flanged male orangutans at Tuanan decreased their long-call rates during and/or after prolonged exposure to wildfire smoke (Erb et al., 2023). In another study of Tuanan orangutans, Ashbury et al. (2022) documented changes in the social behaviour of adult females during low-fruit periods before and after the 2015 fires. Post-fire, they documented decreased probabilities of association between females and weaned immature offspring, and between related and unrelated adult females, and increased probability of agonism.

#### Health and physical condition

Five studies examined the effects of fire and smoke on the condition and health of macaques and orangutans in Borneo. After the drought and fires in 1982-83, long-tailed macaques in Kutai National Park appeared to lose weight and looked emaciated (Berenstain, 1986). At Tuanan, Erb et al. (2018, Erb et al. 2023) used urinalysis to monitor changes in flanged male orangutan energy status, and recorded long calls, before, during, and after smoky periods caused by the 2015 fires. They documented an increase in the proportion of urine samples containing ketones (indicating fat catabolism) as well as reduced vocal quality (i.e., increased harshness, perturbations, and biphonation) in the 3 months following the smoky period compared to the pre-smoke period. At Sebangau, Gwynn et al. (2024) also studied orangutans before, during, and after the smoke in 2015, investigating the prevalence and intensity of gastrointestinal parasites. They found that, following exposure to wildfire smoke, orangutans had a higher prevalence of *Enterbius* and *Trichuris* spp., and greater hookworm infection intensity compared to the pre-smoke period. In a study of orangutan translocations between 2007-2017 across Kalimantan, Sherman et al. (2020) showed that translocations were highest in 2015 and 2016, following the 2015 fires. High levels of translocations occurred in areas affected by fires and adjacent human-modified areas. Of the translocated individuals, approximately half of the starving, malnourished, or underweight orangutans had been driven out of their habitats by the fires in 2015.

## Discussion

Our systematic review highlights the varied impacts of El Niño-driven environmental changes on primates in Southeast Asia, with the majority of studies showing dramatic effects at both population and individual levels. Drought and fire can lead to habitat loss and degradation, as well as extended periods of low fruit production. This can cause dietary, behavioural, and ranging changes in primates, which, in turn, can affect physical condition. Studies show that, in the short term, individuals inhabiting the worst-hit areas may suffer direct mortality due to fire or be displaced from their habitat, leading to human intervention when animals enter human-dominated environments. Studies also show that longer-term consequences include crowding of primates within forest fragments, lower reproductive rates, reduced infant survival rates, and, in rare cases, local extirpation. Smoke inhalation can lead to behavioural changes, such as energy-conserving activity patterns and alterations in vocal behaviour and quality, and may contribute to a decline in overall health (Fig. 1).

Yet, the published literature is limited in both scale and scope. We found only 15 studies on this subject, with narrow geographic and taxonomic representation. Of the 105 primate species inhabiting Southeast Asia (IUCN, 2024), only seven (7%) have been studied (33% of great apes, 16% of small apes, 5% of monkeys, and 0% of tarsiers or strepsirrhines). Whereas orangutans were the most frequently studied taxon, there were no studies on lorises and tarsiers, which together represent 22% of Southeast Asia’s primate species. This limited taxonomic scope reflects a broader bias in primate research—just under half of all recognised primate taxa have been studied, with fieldwork heavily focused on a small number of species at a handful of long-term field stations (Bezanson & McNamara, 2019). This bias leaves substantial gaps in understanding how El Niño impacts the region’s primate diversity.

Though Southeast Asia comprises 11 countries, only one country (Indonesia) and two islands (Borneo and Sumatra) were represented in our literature review. This likely reflects the severity of El Niño effects in these areas compared to other parts of Southeast Asia (Hariadi et al., 2021; Fig. 2). However, several other regions in southern Southeast Asia—such as Java and Sulawesi in Indonesia, Peninsular Malaysia and Malaysian Borneo (ie., Sabah and Sarawak)—are also home to diverse primate communities, yet El Niño’s impact has thus far been unexplored. However, the focus on Sumatra and Indonesian Borneo may be due to the fact that these regions experience the most severe El Niño-related fires in Southeast Asia, driven by both their geographic location and anthropogenic pressure (Field et al., 2016).

The literature predominantly focuses on strong El Niño events, particularly those occurring in 1997-98 and 2014-16, revealing a research bias towards strong events and an increase in research interest in recent years, with a notable absence of research prior to the 1982-83 strong El Niño event. Widespread fires and haze are more likely to occur during strong El Niño events; however, the lack of studies exploring the impacts of moderate events highlights a gap in our understanding of how these more frequent but less severe climatic events might impact primates. Notably, the one study that focused on a moderate El Niño event did report changes in primate behaviour (Cheyne, 2008), suggesting that even less severe events can have consequences for primates.

Fire emerged as the most commonly studied El Niño-driven factor, likely because of its immediate and observable consequences. Of the ten studies that considered fire as a key factor, all but one documented negative impacts on primates, including habitat loss, displacement, and reduced food availability. The exception found that El Niño-driven fires affected only a small proportion of proboscis monkey habitat—likely due to their preference for riverine and coastal habitats, which are more resistant to fire (Toulec et al., 2022).

Smoke also emerged as a significant factor influencing primates. Research thus far has focused on only two species—the Bornean orangutan and the white-bearded gibbon—where smoke exposure was linked to negative effects on health and behaviour. These findings align with evidence from multiple studies on captive macaques exposed to wildfire smoke in the USA underscoring the potential for long-term health consequences, showing increased pregnancy loss and severe health and cognition outcomes for neonates and infants (Capitanio et al., 2022; Berns & Haertel, 2024). Further, a recent global review of wildfire smoke and impacts on wildlife highlighted the breadth of its impacts across species, including respiratory problems, reduced mobility, and heightened vulnerability to disease (Sanderfoot et al., 2022).

The broader range of environmental changes caused by El Niño—such as drought and heat—remain relatively understudied in Southeast Asia’s primates. The few studies that considered these factors found that drought reduces plant productivity, leading to lower fruit availability, which is linked to behavioural and dietary changes and reduced physiological condition (Berenstain, 1986; Mcconkey et al., 2003; Ashbury et al., 2022).

Taken together, the limited environmental, taxonomic, and geographic scope of the studies included in our review likely skews our understanding of how primates are affected across the region. In areas where fire risk is lower, the impacts of El Niño may appear through less visible but still impactful changes, such as shifts in food availability or water sources. While most studies attributed observed changes to fire or smoke, extreme heat and drought can also influence primate behaviour and health directly and indirectly, through changes in habitat quality and food resources driven by altered plant productivity (Berenstain, 1986; Mcconkey et al., 2003). These interconnected factors—rainfall, temperature, fire, and smoke—are rarely considered together, limiting our ability to pinpoint the explicit drivers of primate responses during these climatic events. As a result, this more nuanced understanding of El Niño’s multifaceted impacts remains a critical knowledge gap, constraining our ability to generalise the effects of El Niño across Southeast Asia’s diverse primate populations.

Our review reveals key limitations in our understanding of the long-term impacts of El Niño on primates. A third of the research relied on short-term (≤ 1 year) research (Mcconkey et al., 2003; Cheyne, 2008; Erb et al., 2018, Erb et al., 2023; Gwynn et al., 2024) and/or one-off post-event population surveys (Setiawan et al., 2009; Cattau et al., 2015), which provide valuable snapshots into how primates respond in the immediate aftermath of environmental disturbances. However, they fall short of providing insights into how populations adapt or recover over time. This is particularly important given that the ecological effects of El Niño-driven changes can continue over years or even decades. Furthermore, one-off population surveys—conducted after one or more El Niño events—are informative but limited in their ability to infer causal links between El Niño-driven environmental changes and observed population changes. In contrast, repeated population surveys conducted at various times across a longer period offer a more in-depth understanding of population dynamics in relation to El Niño, allowing a more robust analysis of the impacts of environmental changes on primates.

The few longitudinal studies included in our review (spanning up to 40 years) underscore the importance of continuous monitoring to reveal both short- and long-term impacts. Longitudinal research offers crucial insights into primate resilience and recovery. For instance, two studies presented evidence suggesting that primates are capable of recovering and utilising previously burnt forest within two decades if it is left to regenerate naturally (Russon et al., 2015; Lappan et al., 2021), crucial information that cannot be obtained through shorter research periods. However, such datasets are rare due to practical difficulties in maintaining extended research programs in the field. This, along with the unpredictable nature of El Niño, explains why most research is concentrated at established long-term research sites, where researchers are able to respond opportunistically to environmental changes.

Our review underscores the significant impacts of El Niño on primates and highlights a need for more comprehensive research on primate responses to environmental extremes. Given the likely increase in the frequency and intensity of El Niño due to climate change, there is a pressing need to understand how these events impact primates in areas most vulnerable, particularly Southeast Asia. Given their large size, charismatic appearance, and critical role in ecosystem health and function, primates are an ideal focal group for research on climate-driven events. We argue that more research is needed to expand geographic and taxonomic representation to enhance our understanding of the diversity and severity of El Niño’s impacts across taxa and regions. Utilising advanced research technologies like camera traps, drones, and autonomous recording units could enhance data collection efforts by simultaneously reducing human risk and providing broader-scale insights into population- and species-specific responses to environmental changes. Finally, collaboration is essential, and building a network of researchers, particularly those with long-term study sites, would support the collection of robust, long-term, comparative datasets, strengthening conservation efforts across the region. Considering the potential implications of climate-driven events like El Niño for biodiversity and ecosystem resilience, it is imperative that scientists and conservation practitioners act collectively to address this globally important issue.

## Author contributions

Conceptualisation: EJB; literature review: EJB, WME; writing: EJB, WME; revisions and editing: EJB, WME; visualisation: EJB, WME.

## Acknowledgements

This research received no specific grant from any funding agency, or commercial or not-for-profit sectors.

## Ethical standards

No specific approval was required to conduct this research.

## Data availability

All data curated and analysed for this study are provided in the supplementary material.

## References

Ashbury, A.M., Meric de Bellefon, J., Kunz, J.A., Abdullah, M., Marzec, A.M., Fryns, C., et al. (2022) After the smoke has cleared: Extended low fruit productivity following forest fires decreased gregariousness and social tolerance among wild female Bornean orangutans (Pongo pygmaeus wurmbii). International Journal of Primatology, 43, 189–215.

Berenstain, L. (1986) Responses of Long-Tailed Macaques to Drought and Fire in Eastern Borneo: A Preliminary Report. Biotropica, 18, 257–262.

Bernard, A.B. & Marshall, A.J. (2020) Assessing the state of knowledge of contemporary climate change and primates. Evolutionary Anthropology: Issues, News, and Reviews, 29, 317–331.

Berns, K. & Haertel, A.J. (2024) Excess prenatal loss and respiratory illnesses of infant macaques living outdoors and exposed to wildfire smoke. American Journal of Primatology, 86, e23605.

Bezanson, M. & McNamara, A. (2019) The what and where of primate field research may be failing primate conservation. Evolutionary Anthropology: Issues, News, and Reviews, 28, 166–178.

Capitanio, J.P., Del Rosso, L.A., Gee, N. & Lasley, B.L. (2022) Adverse biobehavioral effects in infants resulting from pregnant rhesus macaques’ exposure to wildfire smoke. Nature Communications, 13, 1774.

Cattau, M.E., Husson, S. & Cheyne, S.M. (2015) Population status of the Bornean orang-utan Pongo pygmaeus in a vanishing forest in Indonesia: the former Mega Rice Project. Oryx, 49, 473–480.

Cheyne, S.M. (2008) Effects of meteorology, astronomical variables, location and human disturbance on the singing apes: Hylobates albibarbis. American Journal of Primatology, 70, 386–392.

Cheyne, S.M., Capilla, B.R.K.A., Supiansyah, Adul, Cahyaningrum, E. & Smith, D.E. (2019) Home range variation and site fidelity of Bornean southern gibbons [Hylobates albibarbis] from 2010–2018. PLOS ONE, 14, e0217784.

Climate Prediction Center Internet Team (2012) Warm (El Nino/Southern Oscillation - ENSO) Episodes in the Tropical Pacific. NOAA /National Weather Service. https://www.cpc.ncep.noaa.gov/products/analysis_monitoring/impacts/warm_IMPACTS.shtml [accessed 17 January 2023].

Climate Prediction Center Internet Team (2024) Historical El Nino /La Nina episodes (1950 - present). NOAA /National Weather Service. https://origin.cpc.ncep.noaa.gov/products/analysis_monitoring/ensostuff/ONI_v5.php [accessed 29 May 2024].

Cochrane, M.A. (2003) Fire science for rainforests. Nature, 421, 913–919. Nature Publishing Group.

Davies, S.J. & Unam, L. (1999) Smoke-haze from the 1997 Indonesian forest fires: effects on pollution levels, local climate, atmospheric CO_2_ concentrations, and tree photosynthesis. Forest Ecology and Management, 124, 137–144.

Erb, W.M., Barrow, E.J., Hofner, A.N., Lecorchick, J.L., Mitra Setia, T. & Vogel, E.R. (2023) Wildfire smoke linked to vocal changes in wild Bornean orangutans. iScience, 26, 107088.

Erb, W.M., Barrow, E.J., Hofner, A.N., Utami-Atmoko, S.S. & Vogel, E.R. (2018) Wildfire smoke impacts activity and energetics of wild Bornean orangutans. Scientific Reports, 8, 7606.

Esri (2021) ENSO Global Weather Patterns [Web map]. arcgis. https://www.arcgis.com/home/item.html?id=ca78667a40e54ea29edf10419a229837 [accessed 26 October 2024].

Estrada, A., Garber, P.A., Rylands, A.B., Roos, C., Fernandez-Duque, E., Di Fiore, A., et al. (2017) Impending extinction crisis of the world’s primates: Why primates matter. Science Advances, 3, e1600946.

Field, R.D., van der Werf, G.R., Fanin, T., Fetzer, E.J., Fuller, R., Jethva, H., et al. (2016) Indonesian fire activity and smoke pollution in 2015 show persistent nonlinear sensitivity to El NiÑo-induced drought. Proceedings of the National Academy of Sciences, 113, 9204–9209.

Fredriksson, G.M., Wich, S.A., & Trisno (2006) Frugivory in sun bears (Helarctos malayanus) is linked to El NiÑo-related fluctuations in fruiting phenology, East Kalimantan, Indonesia. Biological Journal of the Linnean Society, 89, 489–508.

Graham, T.L., Matthews, H.D. & Turner, S.E. (2016) A Global-Scale Evaluation of Primate Exposure and Vulnerability to Climate Change. International Journal of Primatology, 37, 158–174.

Gwynn, A.L., Morrogh-Bernard, H.C., Thornton, A., Segah, H., Azis, A. & Van Veen, F.J.F. (2024) Gastrointestinal parasites of wild Bornean orang-utans (Pongo pygmaeus) in a habitat affected by wildfire smoke. Global Ecology and Conservation, 55, e03214.

Hariadi, M.H., van der Schrier, G., Steeneveld, G.-J., Sopaheluwakan, A., Tank, A.K., Roberts, M.J., et al. (2021) Evaluation of onset, cessation and seasonal precipitation of the Southeast Asia rainy season in CMIP5 regional climate models and HighResMIP global climate models. International Journal of Climatology, 42, 3007–3024.

Harrison, M.E., Cheyne, S.M., Sulistiyanto, Y. & Rieley, J.O. (2007) Biological effects of smoke from dry-season fires in non-burnt areas of the Sabangau peat swamp forest, Central Kalimantan, Indonesia. In Proceedings of The International Symposium and Workshop on Tropical Peatland pp. 27–29. Yogyakarta.

Harrison, M.E., Deere, N.J., Imron, M.A., Nasir, D., Adul Asti, H.A., et al. (2024) Impacts of fire and prospects for recovery in a tropical peat forest ecosystem. Proceedings of the National Academy of Sciences, 121, e2307216121.

Hein, L., Spadaro, J.V., Ostro, B., Hammer, M., Sumarga, E., Salmayenti, R., et al. (2022) The health impacts of Indonesian peatland fires. Environmental Health, 21, 62.

IUCN (2021) IUCN. The IUCN Red List of Threatened Species. Version 2021-3. https://www.iucnredlist.org [accessed 6 June 2022].

IUCN (2024) IUCN. The IUCN Red List of Threatened Species. Version 2024-2. https://www.iucnredlist.org [accessed 13 September 2024].

Khan, M.F., Hamid, A.H., Rahim, H.A., Maulud, K.N.A., Latif, M.T., Nadzir, M.S.M., et al. (2020) El NiÑo driven haze over the Southern Malaysian Peninsula and Borneo. Science of The Total Environment, 730, 139091.

Khandekar, M.L., Murty, T.S., Scott, D. & Baird, W. (2000) The 1997 E. NiÑo, Indonesian Forest Fires and the Malaysian Smoke Problem: A Deadly Combination of Natural and Man-Made Hazard. In Natural Hazards: State-of-the-Art at the End of the Second Millennium (eds G.A. Papadopoulos, T. Murty, S. Venkatesh & R. Blong), pp. 131–144. Springer Netherlands, Dordrecht.

Lappan, S., Sibarani, M., O’Brien, T.G., Nurcahyo, A., Andayani, N., Rustiati, E.L., et al. (2021) Long[term effects of forest fire on habitat use by siamangs in Southern Sumatra. Animal Conservation, 24, 355–366.

Lee, B.P.Y.-H., Davies, Z.G. & Struebig, M.J. (2017) Smoke pollution disrupted biodiversity during the 2015 El NiÑo fires in Southeast Asia. Environmental Research Letters, 12, 094022. IOP Publishing.

Lestari, D.O., Sutriyono, E. Sabaruddin & Iskandar, I. (2018) Severe Drought Event in Indonesia Following 2015/16 El NiÑo/positive Indian Dipole Events. Journal of Physics: Conference Series, 1011, 012040.

Limin, S.H., Rieley, J.O., Jaya, S. & Gumiri, S. (2006) The impact of forest fires and resultant haze on terrestrial ecosystems and human health in central Kalimantan, Indonesia. Tropics, 15, 321–326.

Liu, Y., Zhang, W., Jiang, F., Chen, H.-C., Jin, F.-F. & Hu, S. (2025) Diverse Timing of El NiÑo Onset Linked to Preconditioned Recharge State and Occurrence of Westerly Wind Bursts. Geophysical Research Letters, 52, e2024GL113668.

Marlier, M.E., DeFries, R.S., Voulgarakis, A., Kinney, P.L., Randerson, J.T., Shindell, D.T., et al. (2013) El NiÑo and health risks from landscape fire emissions in southeast Asia. Nature Climate Change, 3, 131–136.

McAlpine, C.A., Johnson, A., Salazar, A., Syktus, J., Wilson, K., Meijaard, E., et al. (2018) Forest loss and Borneo’s climate. Environmental Research Letters, 13, 044009.

Mcconkey, K.R., Ario, A., Aldy, F. & Chivers, D.J. (2003) Influence of Forest Seasonality on Gibbon Food Choice in the Rain Forests of Barito Ulu, Central Kalimantan. International Journal of Primatology, 24, 19–32.

Myers, N., Mittermeier, R.A., Mittermeier, C.G., da Fonseca, G.A.B. & Kent, J. (2000) Biodiversity hotspots for conservation priorities. Nature, 403, 853–858.

Niedzielski, T. (2014) El NiÑo/Southern Oscillation and Selected Environmental Consequences. Advances in Geophysics, 55, 77–122.

O’Brien, T.G., Kinnaird, M.F., Nurcahyo, A., Prasetyaningrum, M. & Iqbal, M. (2003) Fire, demography and the persistence of siamang (Symphalangus syndactylus: Hylobatidae) in a Sumatran rainforest. Animal Conservation, 6, 115–121.

Page, S.E. & Hooijer, A. (2016) In the line of fire: the peatlands of Southeast Asia. Philosophical Transactions of the Royal Society B: Biological Sciences, 371, 20150176.

Page, S.E., Rieley, J.O. & Banks, C.J. (2011) Global and regional importance of the tropical peatland carbon pool. Global Change Biology, 17, 798–818.

Qian, X., Qiu, B. & Zhang, Y. (2019) Widespread Decline in Vegetation Photosynthesis in Southeast Asia Due to the Prolonged Drought During the 2015/2016 El NiÑo. Remote Sensing, 11, 910. Multidisciplinary Digital Publishing Institute.

Ramakreshnan, L., Aghamohammadi, N., Fong, C.S., Bulgiba, A., Zaki, R.A., Wong, L.P. & Sulaiman, N.M. (2018) Haze and health impacts in ASEAN countries: a systematic review. Environmental Science and Pollution Research, 25, 2096–2111.

Rifai, S.W., Li, S. & Malhi, Y. (2019) Coupling of El NiÑo events and long-term warming leads to pervasive climate extremes in the terrestrial tropics. Environmental Research Letters, 14, 105002.

Roy, B. & Goh, N. (2019) A review on smoke haze in Southeast Asia: Deadly impact on health and economy. Quest International Journal of Medical and Health Sciences, 2, 23–28.

Russon, A.E., Kuncoro, P. & Ferisa, A. (2015) Orangutan behavior in Kutai National Park after drought and fire damage: Adjustments to short- and long-term natural forest regeneration. American Journal of Primatology, 77, 1276–1289.

Sanderfoot, O.V., Bassing, S.B., Brusa, J.L., Emmet, R.L., Gillman, S.J., Swift, K. & Gardner, B. (2022) A review of the effects of wildfire smoke on the health and behavior of wildlife. Environmental Research Letters, 16, 123003.

Santoso, A., Mcphaden, M.J. & Cai, W. (2017) The Defining Characteristics of ENSO Extremes and the Strong 2015/2016 El NiÑo. Reviews of Geophysics, 55, 1079–1129.

Setiawan, A., Nugroho, T.S. Djuwantoko & Pudyatmoko, S. (2009) A Survey of Miller’s Grizzled Surili, Presbytis Hosei Canicrus, in East Kalimantan, Indonesia. Primate Conservation, 24, 139–143.

Sherman, J., Ancrenaz, M. & Meijaard, E. (2020) Shifting apes: Conservation and welfare outcomes of Bornean orangutan rescue and release in Kalimantan, Indonesia. Journal for Nature Conservation, 55, 125807.

Shokoohinia, P., Assareh, N., Manomaiphiboon, K., Chusai, C., Kerkkaiwan, S., Unapumnuk, K. & Aman, N. (2020) Impacts of transboundary smoke haze from biomass burning in lower Southeast Asia on air quality in Southern Thailand. Journal of Sustainable Energy & Environment, 11, 1–10.

Sodhi, N.S., Koh, L.P., Brook, B.W. & Ng, P.K.L. (2004) Southeast Asian biodiversity: an impending disaster. Trends in Ecology & Evolution, 19, 654–660.

Sodhi, N.S., Posa, M.R.C., Lee, T.M., Bickford, D., Koh, L.P. & Brook, B.W. (2010) The state and conservation of Southeast Asian biodiversity. Biodiversity and Conservation, 19, 317–328.

Thirumalai, K., DiNezio, P.N., Okumura, Y. & Deser, C. (2017) Extreme temperatures in Southeast Asia caused by El NiÑo and worsened by global warming. Nature Communications, 8, 15531.

Toulec, T., Lhota, S., Scott, K., Putera, A.K.S., Kustiawan, W. & Nijman, V. (2022) A decade of proboscis monkey (Nasalis larvatus) population monitoring in Balikpapan Bay: Confronting predictions with empirical data. American Journal of Primatology, 84, e23357.

Venkatappa, M., Sasaki, N., Han, P. & Abe, I. (2021) Impacts of droughts and floods on croplands and crop production in Southeast Asia – An application of Google Earth Engine. Science of The Total Environment, 795, 148829.

Vos, R. (1999) How to Measure the Cost of Natural Disasters? The Case of ‘El NiÑo’ in Ecuador, 1997–8. Revista Europea de Estudios Latinoamericanos y del Caribe /European Review of Latin American and Caribbean Studies, 21–33.

Wang, B., Luo, X., Yang, Y.-M., Sun, W., Cane, M.A., Cai, W., et al. (2019) Historical change of El NiÑo properties sheds light on future changes of extreme El NiÑo. Proceedings of the National Academy of Sciences, 116, 22512–22517.

Wooster, M.J., Perry, G.L.W. & Zoumas, A. (2012) Fire, drought and El NiÑo relationships on Borneo (Southeast Asia) in the pre-MODIS era (1980–2000). Biogeosciences, 9, 317–340.

Zhang, L., Ameca, E.I., Cowlishaw, G., Pettorelli, N., Foden, W. & Mace, G.M. (2019) Global assessment of primate vulnerability to extreme climatic events. Nature Climate Change, 9, 554–561.

Zhou, T., Wu, B. & Dong, L. (2014) Advances in research of ENSO changes and the associated impacts on Asian-Pacific climate. Asia-Pacific Journal of Atmospheric Sciences, 50, 405–422.

