## Supplemental Material 1 for "A Systematic Review of the Impacts of El Niño-Driven Drought, Fire, and Smoke on Non-Human Primates in Southeast Asia"

Supplementary Table 1 List of moderate or strong El Niño events since 1950-2019. Data were obtained from the National Oceanic and Atmospheric Administration (Climate Prediction Center Internet Team, 2022) which gives a monthly breakdown of ENSO patterns from 1950 to present. Intensity classifications are based on research by Wang et al (2019).

| **Year** | **Strength** |
| --- | --- |
| 1951-52 | Moderate |
| 1953-54 | Moderate |
| 1957-58 | Moderate |
| 1963-64 | Moderate |
| 1965-66 | Moderate |
| 1972-73 | Strong |
| 1976-77 | Moderate |
| 1977-78 | Moderate |
| 1979-80 | Moderate |
| 1982-83 | Strong |
| 1986-88 | Moderate |
| 1991-92 | Moderate |
| 1994-95 | Moderate |
| 1997-98 | Strong |
| 2002-03 | Moderate |
| 2004-05 | Moderate |
| 2006-07 | Moderate |
| 2009-10 | Moderate |
| 2014-16 | Strong |
| 2018-19 | Moderate |

supplementary fig. 1 A schematic diagram illustrating the systematic search steps taken to identify literature to include in this review article. Our initial keyword search yielded 376 articles. After filtering by article type, removing duplicates, and reviewing titles and abstracts, 32 articles underwent a full text review by both authors, resulting in 15 final articles included in the review.

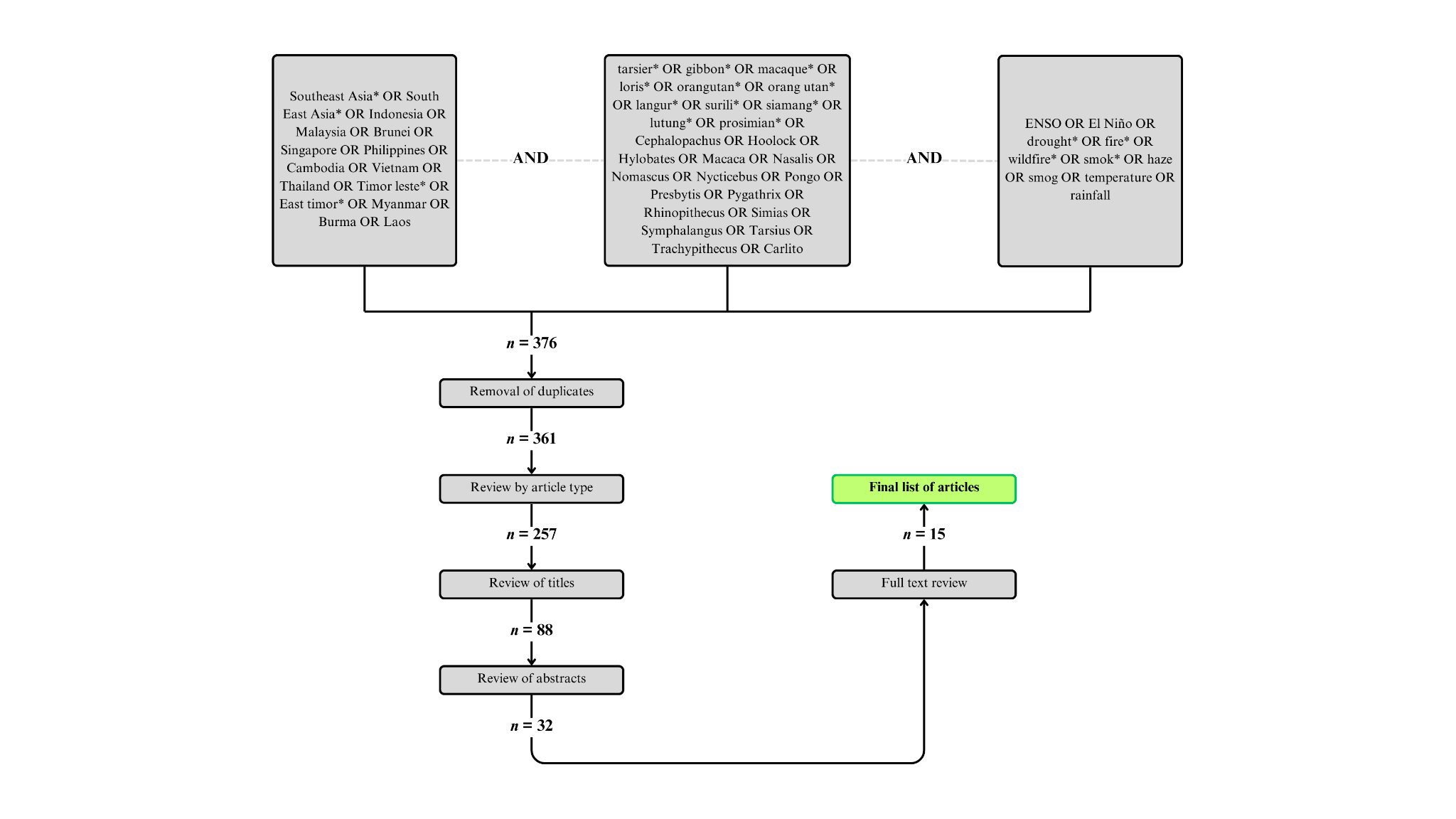

Supplementary Material 1 Search history details downloaded from Web of Science on 2025-04-30.

\# Web of Science Search Strategy (v0.1)

\# Database: All Databases

\# Entitlements:

- WOS: 1900 to 2025

- BCI: 1926 to 2025

- BIOSIS: 1926 to 2025

- CABI: 1910 to 2025

- DRCI: 1900 to 2025

- DIIDW: 1966 to 2025

- FSTA: 1969 to 2025

- GRANTS: 1953 to 2025

- KJD: 1980 to 2025

- MEDLINE: 1950 to 2025

- PCI: 1950 to 2025

- PPRN: 1991 to 2025

- PQDT: 1637 to 2025

- SCIELO: 2002 to 2025

- ZOOREC: 1864 to 2025

\# Searches:

1: ((((((((((((((TS=(Southeast Asia*)) OR TS=(South East Asia*)) OR TS=(Indonesia)) OR TS=(Malaysia)) OR TS=(Brunei)) OR TS=(Singapore)) OR TS=(Philippines)) OR TS=(Cambodia)) OR TS=(Vietnam)) OR TS=(Thailand)) OR TS=(Timor leste*)) OR TS=(East timor*)) OR TS=(Myanmar)) OR TS=(Burma)) OR TS=(Laos) and Preprint Citation Index (Exclude – Database)	Date Run: Wed Apr 30 2025 13:47:46 GMT-0400 (Eastern Daylight Time)	Results: 1347457

2: (((((((((((((((((((((((((((TS=(tarsier*)) OR TS=(gibbon*)) OR TS=(macaque*)) OR TS=(loris*)) OR TS=(orangutan*)) OR TS=(orang utan*)) OR TS=(langur*)) OR TS=(surili*)) OR TS=(siamang*)) OR TS=(lutung*)) OR TS=(prosimian*)) OR TS=(Cephalopachus)) OR TS=(Hoolock)) OR TS=(Hylobates)) OR TS=(Macaca)) OR TS=(Nasalis)) OR TS=(Nomascus)) OR TS=(Nycticebus)) OR TS=(Pongo)) OR TS=(Presbytis)) OR TS=(Pygathrix)) OR TS=(Rhinopithecus)) OR TS=(Simias)) OR TS=(Symphalangus)) OR TS=(Tarsius)) OR TS=(Trachypithecus)) OR TS=(Carlito)) and Preprint Citation Index (Exclude – Database)	Date Run: Wed Apr 30 2025 13:48:32 GMT-0400 (Eastern Daylight Time)	Results: 237296

3: (((((((((TS=(ENSO)) OR TS=(El Niño)) OR TS=(drought*)) OR TS=(fire*)) OR TS=(wildfire*)) OR TS=(smok*)) OR TS=(haze)) OR (smog)) OR TS=(rainfall)) OR TS=(temperature)) and Preprint Citation Index (Exclude – Database)	Date Run: Wed Apr 30 2025 13:49:33 GMT-0400 (Eastern Daylight Time)	Results: 15860145

4: #3 AND #2 AND #1 and Preprint Citation Index (Exclude – Database)	Date Run: Wed Apr 30 2025 13:50:15 GMT-0400 (Eastern Daylight Time)	Results: 376

Supplementary Material 2 Full set of records produced by Web of Science search completed on 2025-04-30 using the search terms and databases described in Supplementary Material 1.
